# Geometric and demographic effects explain contrasting fragmentation-biodiversity relationships across scales

**DOI:** 10.1101/2024.02.01.577731

**Authors:** Stav Gelber, Shane A. Blowes, Jonathan M. Chase, Andreas Huth, Frank M. Schurr, Britta Tietjen, Julian W. Zeller, Felix May

## Abstract

There is consensus that habitat loss is a major driver of biodiversity loss, while the effects of fragmentation, given a constant total habitat amount, are still debated. Here, we use a process-based metacommunity model to show how strongly scale- and context-dependent fragmentation-biodiversity relationships can emerge from the interplay of two types of fragmentation effects - geometric and demographic. Geometric effects arise from the spatial distribution of species and landscape modification, whereas demographic effects reflect long-term changes in species demographic rates following landscape modification. We introduce a novel approach to partitioning these two types of effects and assess how key ecological processes and factors, such as dispersal, habitat heterogeneity, and edge effects, influence geometric, demographic, and net fragmentation effects across spatial scales. We conclude that the framework of geometric and demographic effects can reconcile previous apparently conflicting results and hopefully unlock and advance the debate on biodiversity change in modified landscapes.

## Introduction

Land use change is a dominant driver of biodiversity change with often negative effects on ecosystem services (Sala *et al*. 2000; Hasan *et al*. 2020; Jaureguiberry *et al*. 2022). Land use change modifies both the amount of remaining natural habitat as well as its spatial configuration (Fahrig 2003; Fischer & Lindenmayer 2007). In this context, habitat loss is defined as the reduction in the total habitat amount for a given set of focal species, while habitat fragmentation (also called fragmentation *per se*) refers to changes in spatial habitat configuration, such as when the number of habitat fragments increases and the average fragment size decreases while maintaining a constant total habitat amount (Fahrig 2003). While there is broad consensus that habitat loss negatively affects biodiversity (Brooks et al., 2002; IPBES, 2018), the isolated effects of fragmentation (*per se*) on biodiversity are still intensely debated with some arguing that fragmentation effects can often be positive for common measures of biodiversity (Fahrig *et al*. 2019; Fahrig 2020) and others arguing fragmentation effects are most often negative (Fletcher *et al*. 2018). These seemingly contradicting results may - at least partly - arise from inappropriate comparisons of studies conducted at different spatial scales (i.e., sample-, fragment-, and landscape-scale), underlining that cross-scale extrapolations can be misleading, as the identity, relative importance, magnitude, and even direction of effect of biodiversity drivers can vary with spatial scale (Giladi *et al*. 2011; Chase *et al*. 2018; Fahrig *et al*. 2019).

There is now an emerging consensus highlighting two important aspects, namely the scale- and context-dependence of fragmentation effects on biodiversity (Tscharntke *et al*. 2012; Fahrig *et al*. 2019; Riva & Fahrig 2023). Fragmentation effects have been studied across a large range of spatial scales ranging from comparisons of effort-standardised local samples (alpha diversity) to comparisons of entire landscapes)(gamma diversity). At the local scale, there is consistent evidence that smaller fragments tend to have lower alpha diversity (Haddad et al. 2015; Chase et al. 2020). In contrast, comparisons at the landscape scale often show either neutral or positive relationships between fragmentation and gamma diversity (Quinn & Harrison 1988; Fahrig 2003, 2017, 2020; Riva & Fahrig 2023).

Furthermore, at a given scale, fragmentation effects might be strongly context-dependent as a result of the multiple processes that can affect biodiversity. For instance, several small fragments might capture more gamma diversity in landscapes with high habitat heterogeneity, due to high turnover in species composition among the fragments (i.e., high beta diversity). In contrast, in homogenous landscapes with low compositional turnover (low beta diversity), increased mortality rates in small fragments (e.g. due to demographic stochasticity and/or negative edge effects) might result in a decrease of gamma diversity at the landscape-scale (Fahrig *et al*. 2019, 2022). In other ecosystems, processes such as increased food availability at habitat edges or release from competition or predation in small fragments might even increase local scale diversity in fragmented landscapes (Crooks & Soule 1999; Klingbeil & Willig 2009).

The hypotheses that scale- and context-dependence can reconcile contrasting fragmentation-biodiversity relationships have been formulated rather qualitatively and anecdotally (Tscharntke *et al*. 2012; Fahrig *et al*. 2019). What is missing so far is a mechanistic, quantitative understanding of the combinations of specific ecological processes and biodiversity drivers that can result in scale-dependent changes in the magnitude and direction of fragmentation effects. Likewise, despite long lists of candidate processes that potentially drive context-dependent fragmentation effects (Ewers & Didham 2006; Fischer & Lindenmayer 2007; Fahrig *et al*. 2019, 2022), we lack an understanding of how variations of a given ecological process can result in positive, neutral, or negative fragmentation-biodiversity relationships.

We argue that a quantitative understanding of fragmentation-biodiversity relationships across contexts and scales requires an improved conceptual framework. Here, we adopt a recent framework that distinguishes two steps in the response of biodiversity to fragmentation, namely (i) geometric and (ii) demographic fragmentation effects (May *et al*. 2019). In the first step, geometric fragmentation effects act like a “cookie cutter” on species distributions: All individuals in remaining habitat fragments survive, while all individuals in the converted habitat (matrix) are assumed to be dead (Kinzig & Harte 2000; Green & Ostling 2003, He & Hubbell 2011). Accordingly, geometric effects exclusively depend on the geometry of habitat conversion and the spatial distributions of species prior to that conversion (Chisholm *et al*. 2018, May *et al*. 2019).

The second step involves demographic fragmentation effects that emerge due to the effects of landscape modification on species’ demographic rates (i.e., birth, death, dispersal) (Hanski 1999; Ewers & Didham 2006). These “demographic effects”, can emerge from several mechanisms such as increased demographic stochasticity (Lande 1993), Allee effects (Courchamp *et al*. 2008), positive or negative edge effects (Pfeifer *et al*. 2017), shifts in dispersal among fragments (Hanski *et al*. 2013), or changes in biotic interactions (Crooks & Soule, 1999).

Several previous models could have been used to assess the implication of geometric and demographic fragmentation effects, but to our knowledge, previous studies focussed on single scales (e.g. Rybicki & Hanski 2013; Rybicki et al. 2020), ignored demographic fragmentation effects (Chisholm et al. 2018; May et al. 2019), or did not disentangle geometric vs. demographic effects (e.g., Lasky & Keitt 2013; Rybicki & Hanski 2013).

Here, we present a generic process-based modelling framework aimed at comprehensively analysing the influence of geometric and demographic fragmentation effects on biodiversity at different scales and in different contexts. Our dynamic and spatial metacommunity model sheds new light on fragmentation-biodiversity relationships. Specifically, we show how geometric and demographic fragmentation effects depend on essential ecological factors and processes in natural and fragmented landscapes, including habitat heterogeneity, habitat filtering, dispersal, and edge effects, and how this generates scale- and context-dependent biodiversity-fragmentation relationships.

We develop an approach for partitioning geometric and demographic effects from biodiversity time series. This enables us to investigate whether geometric and demographic effects are linked to the same or distinct processes. The combination of this novel conceptual approach with detailed and extensive mechanistic simulations enhances the understanding of biodiversity responses to fragmentation and improves predictions of scale-dependent biodiversity changes in anthropogenically modified landscapes. Using systematic variations of model parameters, we ask: How do geometric and demographic effects interact to determine scale-dependent biodiversity responses to habitat fragmentation? We show strongly scale-dependent associations of biodiversity to fragmentation and that associations between fragmentation and biodiversity at the landscape-scale can be positive, neutral, or negative depending on the relative importance and the strength of geometric vs. demographic effects.

## Methods

### Model Overview

To explore the scale- and context dependence of fragmentation-biodiversity relationships, we developed a spatially-explicit model that simulates community dynamics in fragmented and heterogeneous landscapes. The model is inspired by previous metacommunity models (Mouquet & Loreau 2003; Gravel *et al*. 2006; Rybicki & Hanski 2013) and includes local community dynamics (i.e., reproduction, mortality, and competition) and dispersal among local communities. The simulated landscapes can feature different degrees of habitat heterogeneity. Accordingly, with appropriate parameter variations, the model is able to embrace the metacommunity archetypes of neutral dynamics, species sorting, and mass effects (Leibold *et al*. 2004; Thompson *et al*. 2020).

The model includes two main entities: individuals and habitat cells. Individuals are characterised by their species-specific environmental niche preference, niche breadth, species ID, and location. Habitat cells are characterised by their type (habitat or matrix), environmental value, carrying capacity, edge category, and location (XY-coordinates). For simplicity, the model makes two broad assumptions: (i) Individuals are not able to move (i.e. it describes sessile organisms such as plants), and (ii) individuals are not able to survive in the matrix cells. Time is simulated in the model with discrete steps (100 time steps per simulation). In each time step, four processes take place sequentially: reproduction, dispersal, death, and immigration from the species pool. The simulation is divided into two temporal phases: (i) initialization with an undisturbed continuous landscape lasting 40 time steps and (ii) a phase with a fragmented landscape lasting 60 time steps. Between these phases, landscape modification takes place using the “cookie-cutter” approach (May *et al*. 2019), and all individuals that are located in matrix cells after landscape modification instantly die. Species distribution patterns and biodiversity at different scales emerge from habitat heterogeneity, survival and recruitment probabilities, dispersal rates, and interspecific interactions via competition for space.

The R code used for running the model is available from GitHub (https://github.com/Stavooo/Gelber_etal_2023_frag)

### Model initialization

We initialise the model with 5,000 individuals of 1,000 different species. Each species has a unique value for its niche preference, which is sampled from a uniform distribution between 0-1. Niche breadth is assumed the same for all species. The individuals are distributed randomly in habitat cells with niche values that fall in the range of the species niche preference value +-its niche breadth.

We generate a heterogeneous landscape with a given environmental autocorrelation which is modified once during simulations (see “Simulating habitat fragmentation”). The patterns of niche values and modifications in the landscape are generated stochastically and thus differ between simulation runs. To include edge effects in the model, edge cells are defined as any cell where one or more of its eight neighbouring cells is a matrix cell. Cells located on the edge of the simulation space which are surrounded by habitat cells are not considered edge cells for the purpose of explicit edge effects (see “Model processes“) but inflict implicit negative effects as the simulation space is not torus-shaped and propagules have higher chance to land outside of the simulation landscape.

### Simulating habitat fragmentation

The landscape for simulations is a square 50 × 50 grid containing two types of cells: habitat and matrix cells. Each habitat cell has a pre-defined carrying capacity (50), limiting the number of individuals that can co-exist in the cell. To simulate environmental heterogeneity across the landscape, we assign an environmental value from the interval 0-1 to each habitat cell.

The environmental values are simulated by generating two-dimensional neutral landscapes using fractional Brownian motion (Travis & Dytham 2004; Schlather *et al*. 2015, Sciaini *et al*. 2018). When creating the landscapes, we control the spatial autocorrelation of environmental variation using the Hurst coefficient H, which varies the ‘smoothness’ of the landscape (Hurst 1951). The H values range from 0 (rugged landscape) to 1 (smooth landscape) (Fig. 1).

**Figure 1.**
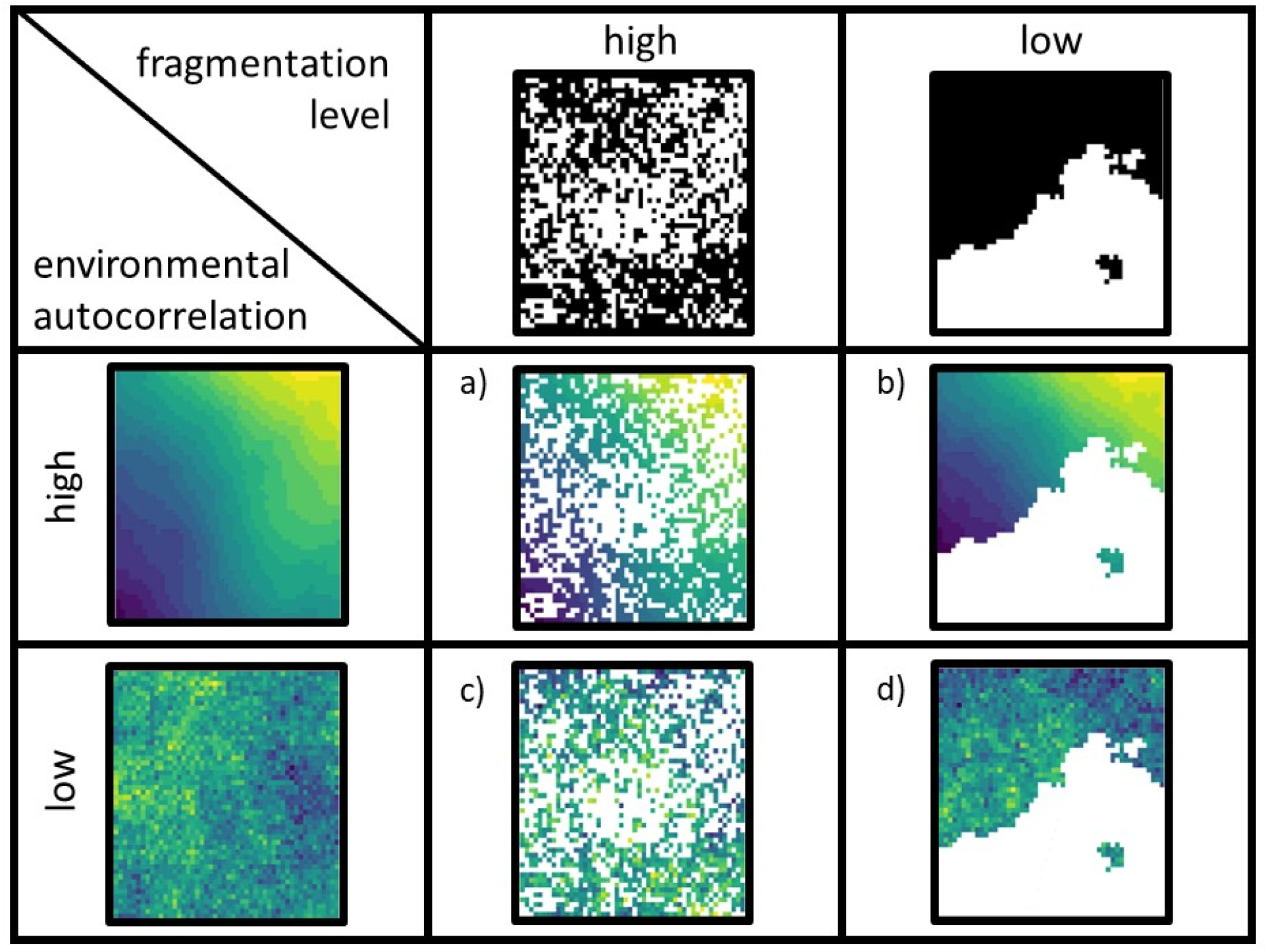
Illustration of the landscape generation process throughout model simulations. The final landscapes (a to d) represent the combination of two landscapes with varying levels of fragmentation and spatial autocorrelation. The habitat amount in the final landscapes is constant. The Hurst coefficient (H) controls for the autocorrelation when generating the continuous landscape (left column) and for the level of fragmentation when generating the spatial pattern of landscape modification (top row). For low autocorrelation and high level of fragmentation, the H value is 0.1, and for high autocorrelation and low fragmentation, the H value is 0.9.

At the start of each simulation, a continuous landscape containing only habitat cells is created with a set degree of spatial environmental autocorrelation (Fig. 1 left column). To modify the landscape following the initialization phase, we create a second and independent landscape, in which we set each cell to be either habitat or matrix by assigning a specific binarization threshold (Fig. 1 top row), which controls the habitat amount in the modified landscape. In this binary landscape, the Hurst coefficient controls the level of fragmentation. A rugged binarized landscape (low H) results in a highly fragmented landscape, while a smooth binarized landscape (high H) results in a landscape with low fragmentation. Finally, to modify the continuous landscape, we replace all habitat cell values in the modified (binarized) landscape with the environmental values from the continuous one (Fig. 1a-d).

### Model Processes

#### Reproduction, dispersal, and establishment

During the reproduction process, the model loops through all individuals, checking for a set of conditions before allowing a reproduction event. In every time step, each individual produces one propagule with a given probability (0.85). To determine the propagule’s target cell, we use an exponential dispersal kernel with a given mean dispersal distance. If the target cell is a habitat cell and its total abundance is below its carrying capacity, the establishment probability of the propagule in the target cell is calculated following Gravel et al. (2006):

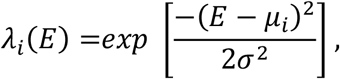

Where λ_*i*_ is the establishment probability of species *i, E* is the cell environmental value, μ_*i*_ is the niche preference value for species *i*, and σ is the niche breadth. Accordingly, the establishment probability equals 1 if the propagule encounters its species-specific environmental optimum and decays exponentially for values lower or higher than the optimum.

#### Death

At each time step the probability that an individual dies is 0.25. In addition, edge effects can alter the death probability of individuals located in edge cells. In this case, the death probability of the individual is multiplied by an edge effect factor ranging between 0.6 (lower mortality, i.e. positive edge effect) and 1.5 (higher mortality, i.e. negative edge effect).

#### Immigration

To simulate the immigration of species from a regional species pool, 50 new individuals are introduced into the landscape in each time step. The species identity of each individual is randomly selected from a species pool of 1,000 species with equal relative abundances of all species. Immigrants are placed in a randomly selected cell and can only be established according to the rules described above (see “Reproduction, dispersal, and establishment”). Since the model does not simulate speciation, species richness would decline over time due to ecological drift without this process of immigration.

### Simulation experiments and model analysis

We examined the scale-dependence of fragmentation effects on biodiversity at two spatial scales. Specifically, species richness was quantified at the landscape-scale (i.e. the whole simulated landscape) and at the sample-scale, which was evaluated at the (habitat) cell scale, and calculated as the average species richness of 30 random habitat cells (Supplementary Material, Fig. S1). Furthermore, landscape-scale and sample-scale species richness were recorded at two time points of every simulation, first directly after landscape modification (step 40) and second at the end of the simulation (step 100) (Fig. 2). These two time points are essential for partitioning geometric and demographic effects (see “Partitioning of geometric and demographic fragmentation effects”). Since this study is focused on fragmentation effects, we used a constant habitat amount of 15% throughout all simulations and varied fragmentation (Hurst coefficient 0.1 - 0.9).

**Figure 2.**
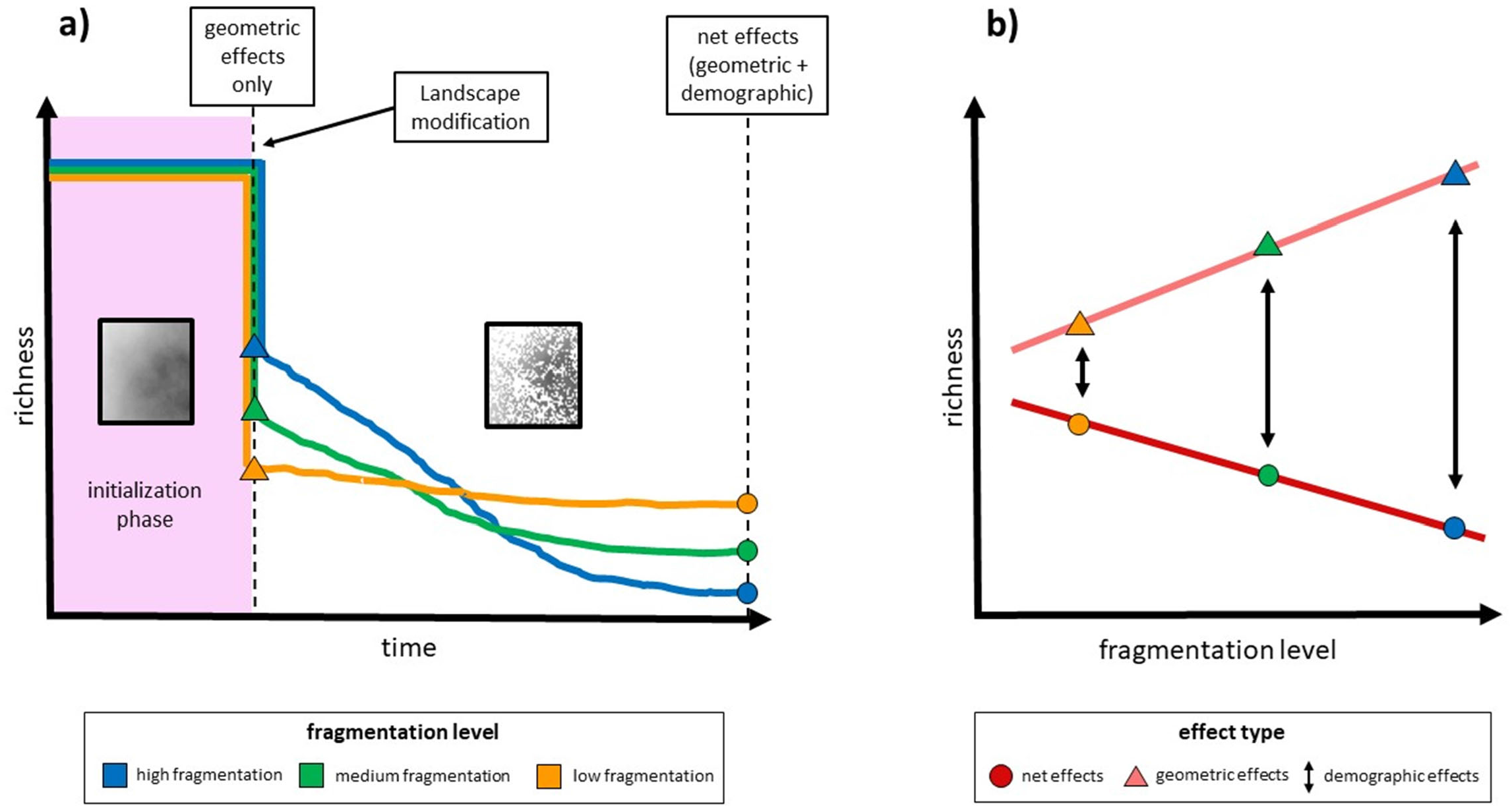
Illustration of the simulation approach to partition geometric and demographic effects based on the model temporal development. Dashed lines represent the time points in which richness data was recorded to expose geometric and net fragmentation effects. The purple arrow indicates the time point in which landscape modification occurred. The three coloured lines represent species richness data for three simulations with varying levels of fragmentation. Grey boxes represent the landscape before and after landscape modification. The values are not taken from simulated data but are an illustration of one possible outcome where geometric fragmentation effects are positive but net effects are negative.

In the first simulation experiment, we used a single reference parameter set (Supplementary Material, Table S2) to illustrate the spatial and temporal scale-dependence of fragmentation-biodiversity relationships.

In the second simulation experiment, we assessed the influence of dispersal distance on fragmentation effects by varying dispersal distance from 1 to 6. May et al. (2019) argued that all processes and factors that determine intraspecific aggregation will also influence geometric fragmentation effects. Therefore, we expect that increasing dispersal will reduce intraspecific aggregation and thus geometric effects. However, with our novel partitioning approach (see “Partitioning of geometric and demographic fragmentation effects”), we can also assess the role of dispersal for demographic fragmentation effects.

To further explore the context-dependence of fragmentation-biodiversity relationships, we varied two other factors in combination with dispersal, specifically environmental autocorrelation and edge effects. Similar to dispersal, increasing environmental autocorrelation should result in higher intraspecific aggregation and stronger geometric effects. It is an open question if autocorrelation also drives demographic effects. In the model, edge effects can change the mortality of individuals located in edge cells. That means, edge effects only influence individuals after landscape modification (since no edges exist in a non-modified landscape), and therefore edge effects by definition can only drive demographic fragmentation effects.

In a third simulation experiment, we simulated a full-factorial design with 10 levels of dispersal distance and 10 levels of the edge effects (0.6 - 1.5), which range from negative to positive edge effects. In a fourth simulation experiment, we simulated a full-factorial design with the same 10 levels of edge effects combined with 10 levels of landscape autocorrelation (0.01 - 0.91).

An overview of all model parameters and their settings is provided in Table S2. All simulation results are based on ten repetitions for every parameter combination. All analyses were performed using R Statistical Software (R Core Team 2022) and simulations were performed on the FU Berlin High-Performance computer, Curta (Bennett *et al*. 2020).

### Partitioning of geometric and demographic fragmentation effects

We applied a novel approach to partition geometric and demographic fragmentation effects to the model simulations that differed in the settings for fragmentation but where all other parameter settings were equal. Intuitively, fragmentation effects on species richness were quantified by fitting simple linear regressions with species richness as the response variable and fragmentation as the predictor variable. In our framework, geometric vs. demographic effects can be disentangled by using species richness derived at different time points of the simulation (Fig. 2).

By definition, the slope of the species richness vs. fragmentation regression measures geometric fragmentation effects, when looking at simulated species richness directly after landscape modification, because, at this time point, all richness differences can exclusively be attributed to the spatial distributions of individuals and habitat, but not to metacommunity dynamics in fragmented landscapes. The challenge is that the simulated richness differences at the end of the simulations include both geometric as well as demographic effects.

We suggest that demographic effects can be isolated in the following way: When there would be no demographic effects the slope of the richness-fragmentation regression should be constant throughout the entire simulation. Thus, any changes in the slope estimate can be attributed to demographic fragmentation effects. Accordingly, we measure demographic effects as the difference between the slope of the species richness-fragmentation regression at the end of the simulation (demographic and geometric effects) and the slope directly after landscape modification (geometric effects only) (Fig. 2). Statistically, this corresponds to the interaction effect between fragmentation and time point. That means negative demographic fragmentation effects will be indicated by decreasing slopes over time (more shallow increase or steeper decrease of species richness with fragmentation), while positive demographic effects will be indicated by increasing slopes over time (steeper increase or more shallow decrease of richness with fragmentation).

## Results

### Scale-dependence of geometric and demographic fragmentation effects

By comparing results at two spatial scales, we explored the scale-dependence of fragmentation effects on biodiversity (Fig. 3). At the sample-scale, geometric effects were neutral across all fragmentation levels by definition, because comparisons of single samples do not include any differences in the geometry of spatial sampling. In contrast, at the landscape-scale, geometric effects were strongly positive, i.e., species richness increased with increasing levels of fragmentation. Identifying the isolated geometric effects provided insights into the mechanisms underlying net fragmentation effects. At the sample-scale, net effects with increasing fragmentation were strongly negative, while at the landscape-scale, net fragmentation effects with increasing fragmentation were weakly positive. Observing the difference between geometric and net effects at each spatial scale revealed the strength of demographic effects. Our simulations thus showed that the apparently contrasting net effects of fragmentation at the sample-scale (negative) and the landscape-scale (neutral to positive) arose from the interplay of immediate geometric and longer-term demographic effects: At the sample-scale, the negative demographic effects dominated, while at the landscape-scale, the positive geometric effects and the negative demographic effects almost cancelled each other out with slight dominance of the geometric effects in this example setting.

**Figure 3.**
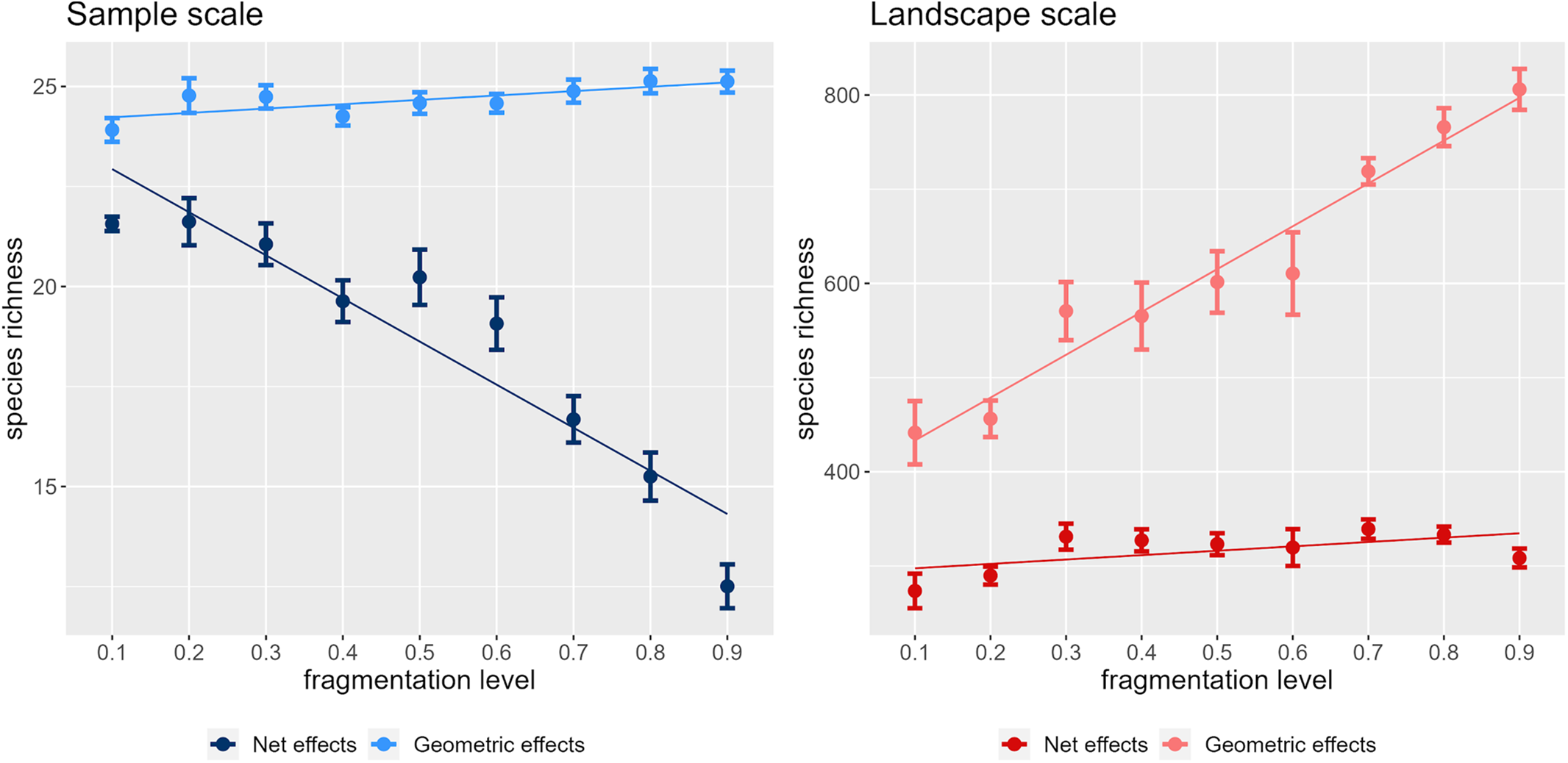
Species richness for the sample scale (blue) and the whole landscape (red) partitioned into geometric effects (light shades), and net effects (dark shades). The results for the landscape and the sample scale are derived from the same simulations. Sample-scale was defined as a mean of 30 randomly-selected habitat cells (left panel), and landscape-scale as the whole simulation space (left panel). Partitioning into geometric and net effects was done by recording results directly after fragmentation for geometric effects and at the end of simulations for net effects. The strength of fragmentation effects on biodiversity is expressed as the slope of a regression line for species richness with increasing levels of fragmentation. Demographic effects are defined as the slope for net effects minus the slope for geometric effects, i.e. demographic effects = net effects – geometric effects. Parameters were set according to the default values (See Table S2).

### Context-dependence of geometric and demographic fragmentation effects

To explore the influence of dispersal distance dynamics on fragmentation effects at the two spatial scales, we ran model simulations with varying dispersal distances (Fig. 4). At the sample-scale, geometric effects were neutral by definition, while negative demographic effects became more pronounced with increasing dispersal distance. As expected, at the landscape-scale, increasing dispersal distances resulted in reduced positive geometric effects as indicated by decreasing slopes of the biodiversity-fragmentation relationships (Fig. 4, bottom). The strength of negative demographic effects increased (i.e., became more negative) from short to intermediate dispersal distance and slightly decreased again with further increases in dispersal, resulting in a weakly unimodal relationship between dispersal distance and demographic effects (Fig. 4, bottom). With increasing dispersal, the combination of reduced positive geometric effects and increased negative demographic effects resulted in a qualitative change from positive to negative net effects of fragmentation at the landscape-scale. That means, just varying a single key ecological process can switch landscape-scale fragmentation-biodiversity relationships from increasing to decreasing, via the joint implications of dispersal on geometric as well as demographic fragmentation effects.

**Figure 4.**
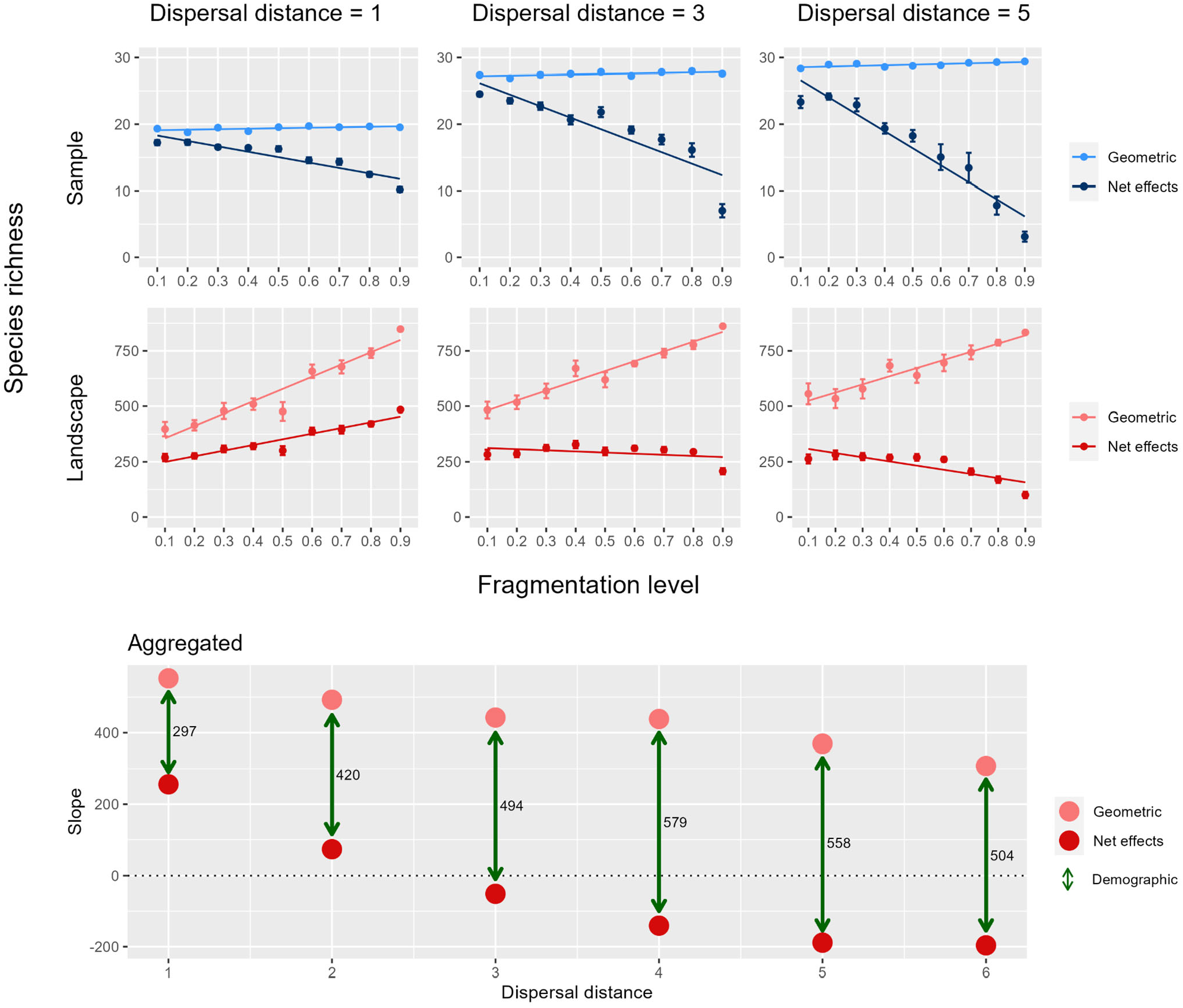
The top six panels show species richness for the whole landscape (red) and for the sample scale (blue) partitioned to geometric effects (light shades), and net effects (dark shades) with three dispersal distances (1, 3, and 5 grid cells). The Bottom panel summarises the results of the top panels including three additional dispersal distances (2, 4, and 6 grid cells). Results are for the landscape scale only and values represent the slope of a regression line for the corresponding parameter setting. Green arrows illustrate the distance between the slope of geometric effects only and net effects, representing the strength of demographic effects in the simulation. The dotted horizontal line indicates where the direction of the net effect changes from positive to negative.

To further explore the context-dependence of fragmentation effects, we assessed the interacting effects of dispersal, edge effects, and environmental autocorrelation. Here, we present the findings for the landscape-scale, while the corresponding simulation results for the sample-scale are provided in the supplementary material (Fig. S3).

When we varied dispersal distance and edge effects simultaneously, geometric effects became weaker with increasing dispersal distance but were not influenced by changing edge effects (Fig. 5a). By definition, edge effects did not influence geometric effects, because edge cells exist only during the simulation phase after landscape modification. Demographic effects were interactively influenced by edge effects and dispersal distance (Fig. 5b). In general, demographic effects were rather weak with positive edge effects and became clearly negative with negative edge effects. The response of demographic effects to dispersal distance appears more complex. Specifically, the relationship between dispersal and negative demographic effects varies from neutral with positive edge effects over unimodal with neutral edge effects (compare Fig. 4 bottom panel, Fig. 5b) to clearly decreasing relationships with negative edge effects. Finally, the interplay between geometric and demographic fragmentation effects resulted in a wide spectrum of net effects on biodiversity, ranging from positive (shorter dispersal and positive edge effects) to negative (longer dispersal and negative edge effects) (Fig. 5c).

**Figure 5.**
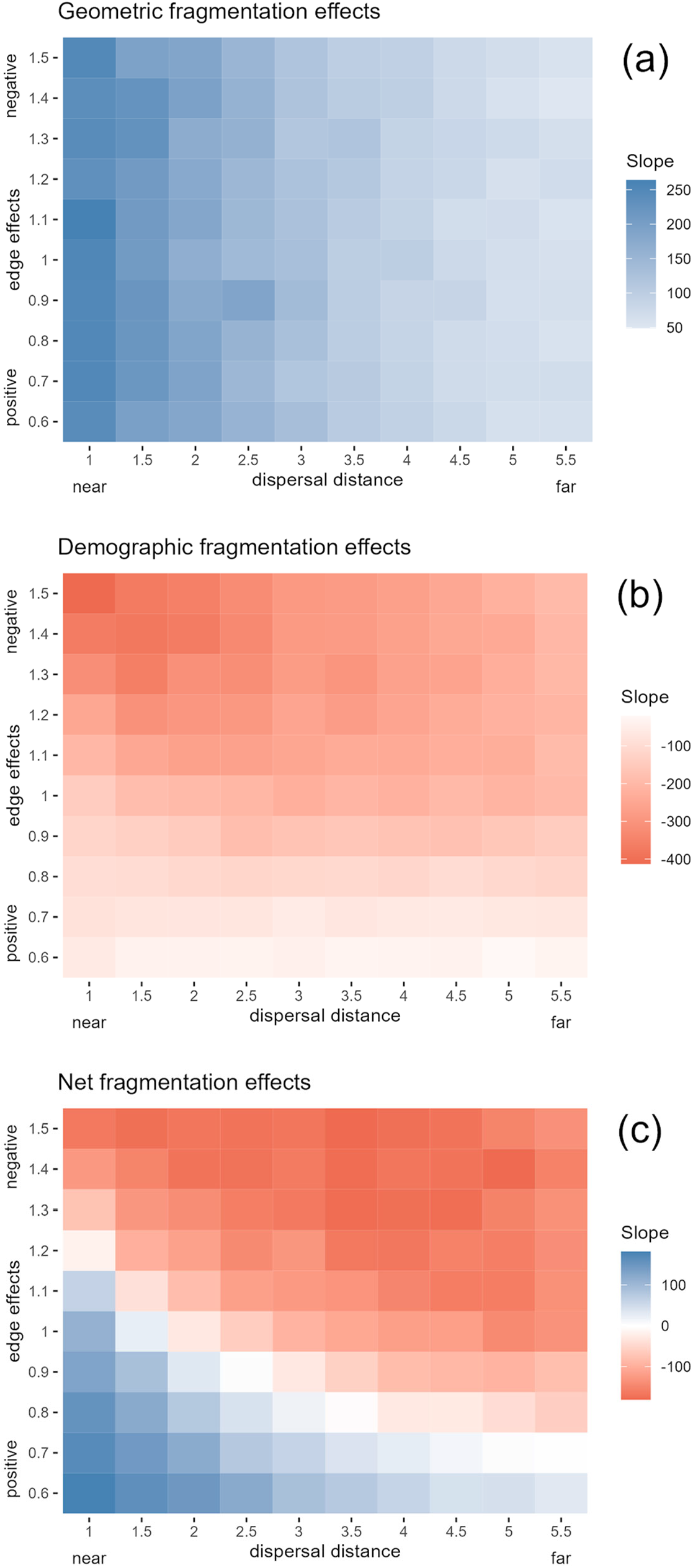
A full factorial design testing the interplay between fragmentation effects at the landscape scale by varying the edge effects and dispersal distance parameters. Plot colours represent the slope of a regression line for species richness with increasing levels of fragmentation (0.1-0.9) immediately after fragmentation for geometric effects (a) and at the end of the simulation for net effects (c). Demographic effects were calculated by deducting the geometric effects slope value from the net effect slope value (b). Slope results for each box were calculated from 9 simulations with varying fragmentation levels repeated 10 times.

We continued to vary landscape autocorrelation and edge effects simultaneously (Fig. 6). As expected, geometric effects increased, i.e., became more positive, with higher landscape autocorrelation (Fig. 6a). Demographic effects were influenced by both landscape autocorrelation and edge effects, where negative demographic effects increased with increasing autocorrelation and negative edge effects (Fig. 6b). Finally, with increasing landscape autocorrelation, net fragmentation effects (Fig. 6c) varied widely from positive to negative. Fig. 6 also clearly indicates a positive correlation between geometric and demographic effects. This is a direct consequence of the partitioning approach used here. With equal net effects of fragmentation, strongly positive geometric effects directly imply strongly negative demographic effects.

**Figure 6.**
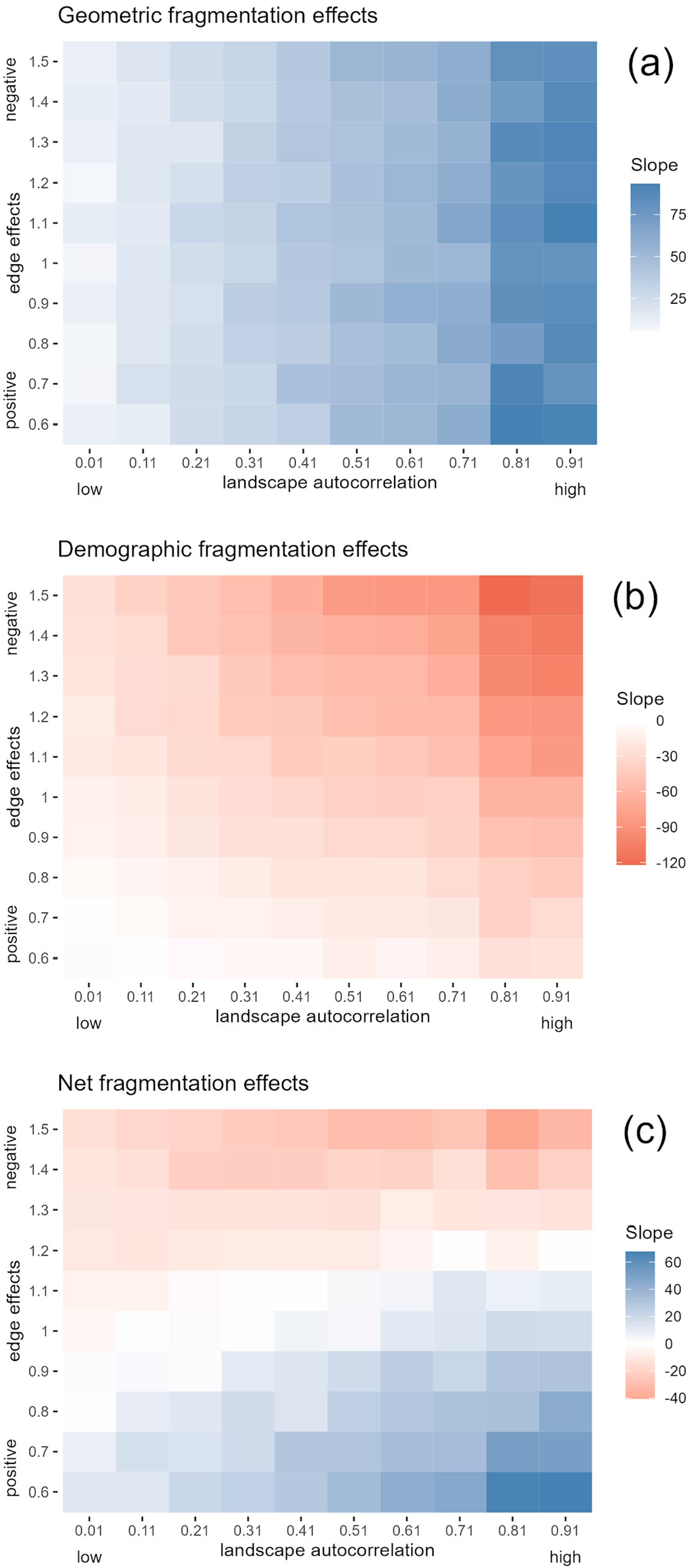
A full factorial design testing the interplay between fragmentation effects at the landscape scale by varying the edge effects and landscape autocorrelation parameters. Plot colours represent the slope of a regression line for species richness with increasing levels of fragmentation. Slope results for each box were calculated from 9 simulations with varying fragmentation levels repeated 10 times.

## Discussion

The effects of fragmentation on biodiversity feature a strong dependence on the spatial scale and the ecological context of studies, but it has remained challenging to incorporate these variations into a single overarching framework (Fletcher *et al*. 2018; Fahrig *et al*. 2019; Valente *et al*. 2023). Here, we have taken a step in this direction by illustrating how scale- and context-dependent fragmentation-biodiversity relationships can be understood and reconciled when we distinguish two types of fragmentation effects – geometric and demographic – and their variation in the context of metacommunity dynamics.

First, we demonstrated both the magnitude as well as the direction of fragmentation effects could change across spatial scales. While scale-dependence in habitat fragmentation effects has been highlighted before (e.g., Tscharntke et al. 2012, Fahrig *et al*. 2019), we were able to show the role that geometric and demographic effects play in the emergence of scale-dependence. We found that positive geometric effects emerge at the landscape-scale because of aggregated species distributions due to local dispersal and/or spatial environmental autocorrelation (Condit *et al*. 2000; McGill 2010). In our model, negative demographic effects emerged even without edge effects as an explicit implementation of demographic effects. This is likely because increasing fragmentation results in a higher proportion of propagules ending up in the matrix and thus increasing dispersal mortality is the main reason for the emerging negative demographic effects in our model. Overall, the interplay between dominant negative demographic effects at the local scale and dominant positive geometric effects at the landscape-scale leads to the disparity between net fragmentation effects across spatial scales. This explanation can reconcile the apparent empirical mismatch between negative fragmentation effects at the sample-scale (Haddad *et al*. 2015; Chase *et al*. 2020) and positive effects at the landscape-scale (Fahrig 2017, Riva & Fahrig 2023).

Second, we explored the context-dependence of net fragmentation effects by varying ecological processes and factors within simulations. As expected, geometric effects were determined by processes and factors that influence species aggregation prior to landscape modification. Accordingly, positive geometric effects decreased with increasing dispersal distance and increased with increasing environmental autocorrelation.

Demographic effects on the other hand were not coupled to a distinct single ecological process and were affected by multiple factors simultaneously. With no or weak edge effects, our simulations showed a unimodal relationship between dispersal distance and the strength of demographic effects, i.e., the strongest negative demographic effects were observed with medium dispersal distance. This finding matches well with the empirical evidence of Thomas (2000), who found that butterfly species with intermediate dispersal distances are more sensitive to fragmentation than species with high or low dispersal rates. He attributed the observed pattern to successful between-patches migration with longer dispersal distances and dispersal within patches with shorter dispersal while intermediate dispersal species were more often dispersed to the matrix. In general, despite the conceptual differentiation between geometric and demographic effects, we found that variations in a single ecological process - dispersal - influence both, geometric as well as demographic effects and thus impose an inherent correlation between the strength of both effects. This highlights why we need a quantitative understanding of which ecological processes and factors influence geometric and demographic effects independently or simultaneously.

As expected, negative edge effects resulted in stronger negative demographic effects and positive edge effects in weaker ones. We found that the emerging demographic effects in the model were never positive (Figs. 5, 6), most likely, because even with positive edge effects the high dispersal mortality in the matrix counteracted these positive demographic effects in high fragmentation scenarios. Accordingly, the high variation of edge effects among species and landscapes (Pfeifer et al. 2017) is likely an additional important driver of the context-dependence of fragmentation-biodiversity relationships.

In our simulations, strongly positive geometric effects tended to be associated with strongly negative edge effects (Figs. 5, 6). We argue that this prediction might be of concern for biodiversity conservation. When a highly fragmented landscape is species-rich directly after landscape modification (i.e., positive geometric effects), most of these species will be rare because of a limited landscape carrying capacity. Therefore, many of these rare species might be at risk of extinction in the long run during metacommunity dynamics in the fragmented landscape (i.e., negative demographic effects).

Recently, Fahrig et al. (2022) suggested the “SLOSS cube hypothesis” to predict in which contexts, we should expect positive, neutral, or negative net fragmentation effects on biodiversity at the landscape-scale. According to this hypothesis, the direction of fragmentation effects primarily depends on three factors: First species clumping (i.e. intraspecific aggregation), second the connectivity among habitat fragments which determines the frequency of species migration between fragments, and third, the role of risk-spreading among several habitat fragments. Our framework includes close relationships to the SLOSS cube but also highlights important extensions. First, our approach embraces different spatial scales, while the SLOSS cube only focuses on the landscape-scale. Second, while the SLOSS cube implies that its three axes can vary independently, our process-based framework challenges this perspective due to the potential joint dependence of the cube’s axes on several processes. While Fahrig et al. (2022) suggested a complete cube with three axes, our process-based perspective predicts real ecosystems will only cover a subspace of the cube due to constraints and correlations arising from ecological processes such as dispersal distance.

A noteworthy difference between how real landscapes are modified and our simulation is that we assume the landscape is modified instantaneously while in reality, it is usually a more gradual process. This assumption is useful for partitioning geometric and demographic effects. Nevertheless, we expect that the key processes we have identified will depend on the species distributions prior to landscape modification (geometric effects) and the response of demographic processes to land use changes (demographic effects) in a manner similar to that which we have outlined. In addition, our model assumes that habitats are lost regardless of their environmental value, while real land use change will often be non-random and target specific habitat types and will depend on factors such as topography (Seabloom *et al*. 2002). The effect of such non-random habitat loss on biodiversity will strongly depend on the quality of the converted habitat type but is unlikely to change the qualitative effects of increasing fragmentation that we observed here. Furthermore, we consider the matrix to be inhospitable to species following landscape modification, whereas modifications could allow for species to persist, at least for some period, in the matrix as is often observed (Jules & Shahani 2003). Finally, several ecological processes can lead to positive demographic effects of fragmentation on biodiversity, such as spread-of-risk, reduced competition, or increased persistence in consumer-resource systems (Simberloff 1988; Fahrig 2017; Pfeifer *et al*. 2017), which would be fruitful to address in future investigations.

Empirical tests of our framework are critical for disentangling geometric and demographic fragmentation effects in real-world ecosystems. Unfortunately, to our knowledge, the spatio-temporal data that would be necessary to apply our partitioning are currently not available because long-term studies of fragmentation tend to focus only on smaller scales (Haddad et al. 2015). It is possible, however, that automated biodiversity monitoring in combination with remote sensing data on land cover at high spatial and temporal resolution (Fischer *et al*. 2021; Ma *et al*. 2023) could provide the data necessary to disentangle geometric and demographic effects in real modified landscapes.

We conclude that partitioning and quantifying geometric vs. demographic fragmentation effects unifies and progresses our understanding of how fragmentation affects biodiversity across spatial scales. We envision that the conceptual framework illustrated here will help to unlock the debate on fragmentation and biodiversity (Valente *et al*. 2023). Despite the simplicity and generality of our approach, our mechanistic simulations provide comprehensive explanations for the strong dependence of fragmentation-biodiversity relationships on spatial scale and ecological context. Still, our results also highlight the potentially complex relationship of ecological processes to geometric and demographic effects. Both types of effects can be influenced by partly overlapping sets of processes and factors, which can have synergistic or antagonistic effects. We suggest that the partitioning of geometric and demographic fragmentation effects in empirical data represents a relevant and needed next step for a deeper understanding of biodiversity change in modified landscapes.

## Acknowledgements

SG acknowledges funding from the German Research Foundation (DFG), grant number MA 5962/1-1. We thank Franziska Taubert for her feedback on the manuscript and Selina Baldauf for reviewing and commenting on the simulation model code.

