## Supporting Information for "Geometric and demographic effects explain contrasting fragmentation-biodiversity relationships across scales"

### S1 Spatial scales in the model

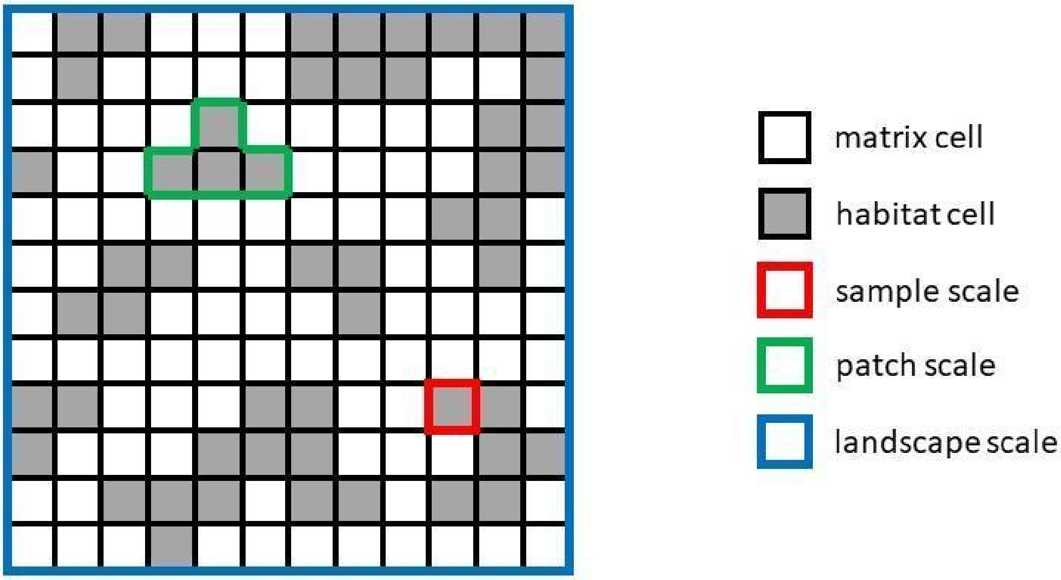


*Figure S1: The figure illustrates how we defined spatial scales within model simulations. We defined "sample-scale" as single cells and "landscape-scale" as the whole simulation space - a square grid with a side length of 50 cells. "Patch-scale" was defined as a clump of neighbouring cells but was not used in the final analysis.*

### S2 Summary table of model parameters and their units

**Table S2**
*Summary table of model parameters and their units*

| Parameter name | Functionality | Accepted values | Default value |
|---|---|---|---|
| Grid side length | Determine simulation grid size | ≥ 1 | 50 |
| Habitat amount | The proportion of habitat amount following fragmentation | 0 - 1 | 0.15 |
| Spatial autocorrelation | Controls for landscape smoothness/roughness | 0 - 1 | 0.9 |
| Fragmentation | Level of fragmentation after modification | 0 - 1 | 0.1-0.9 |
| Population size | The initial number of individuals | ≥ 1 | 5,000 |
| Species | Number of species in the species pool | ≥ 1 | 1,000 |
| Niche breadth | Determine the range for accepted niche values | 0 - 1 | 0.1 |
| Reproduction rate | Proportion of successful reproduction events in each time step | 0 - 1 | 0.85 |
| Death rate | Proportion of death events in each time step | 0 - 1 | 0.25 |
| Dispersal distance | Mean distance for the dispersal kernel | ≥ 1 | 1-6 |
| Carrying capacity | Amount of individuals a cell can host | ≥ 1 | 50 |
| Immigrants | Amount of new individuals introduced at each time step | ≥ 1 | 50 |
| Edge effects | Type and strength of explicit edge effects | 0 - 2 | 0.6-1.5 |

### S3 Heatmaps for sample-scale

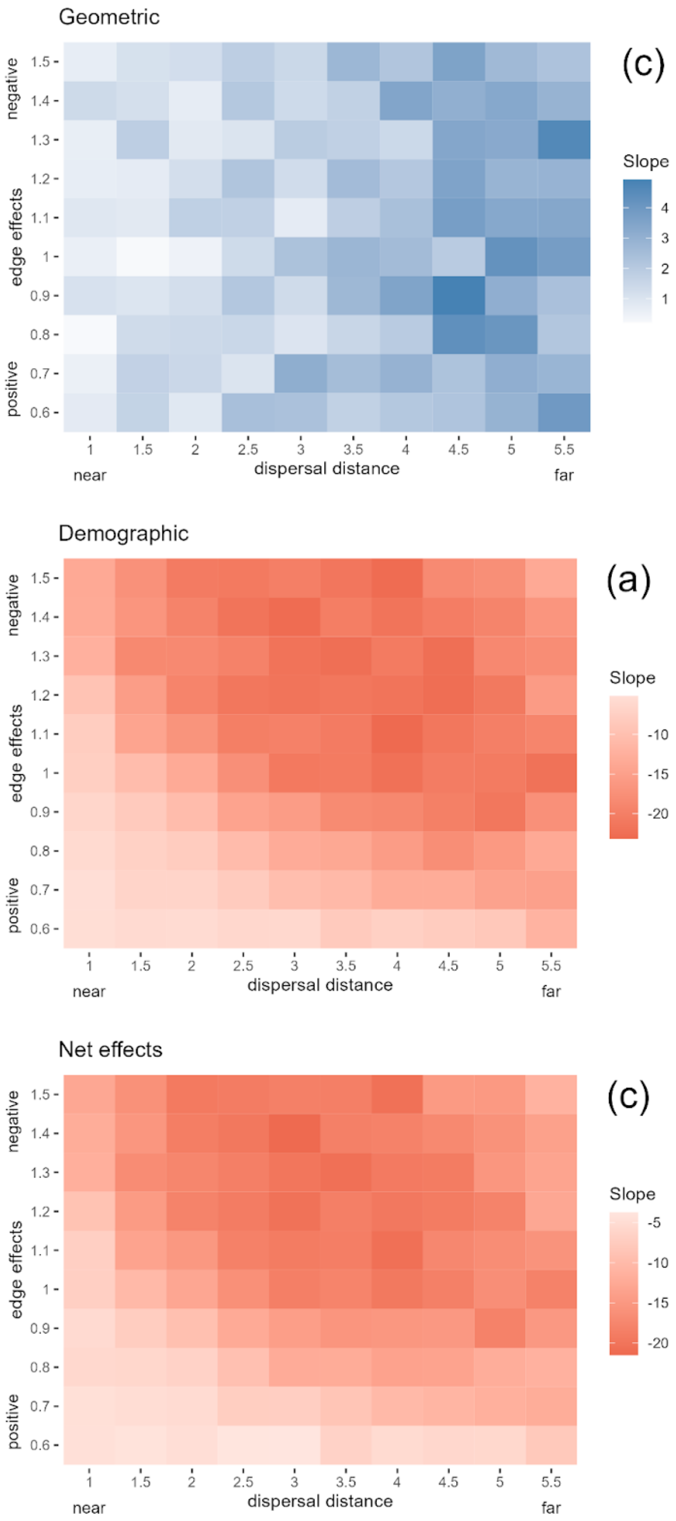


*Figure S3A: A full factorial design testing the interplay between fragmentation effects at the sample scale by varying the edge effects and dispersal distance parameters. Plot colours represent the slope of a regression line for species richness with increasing levels of fragmentation (0.1-0.9) immediately after fragmentation for geometric effects (a) and at the end of the simulation for net effects (c). Demographic effects were calculated by deducting the geometric effects slope value from the net effect slope value (b). Slope results for each box were calculated from 9 simulations with varying fragmentation levels repeated 10 times. Values for species richness at the sample scale are based on mean values of 30 samples (single cells).*

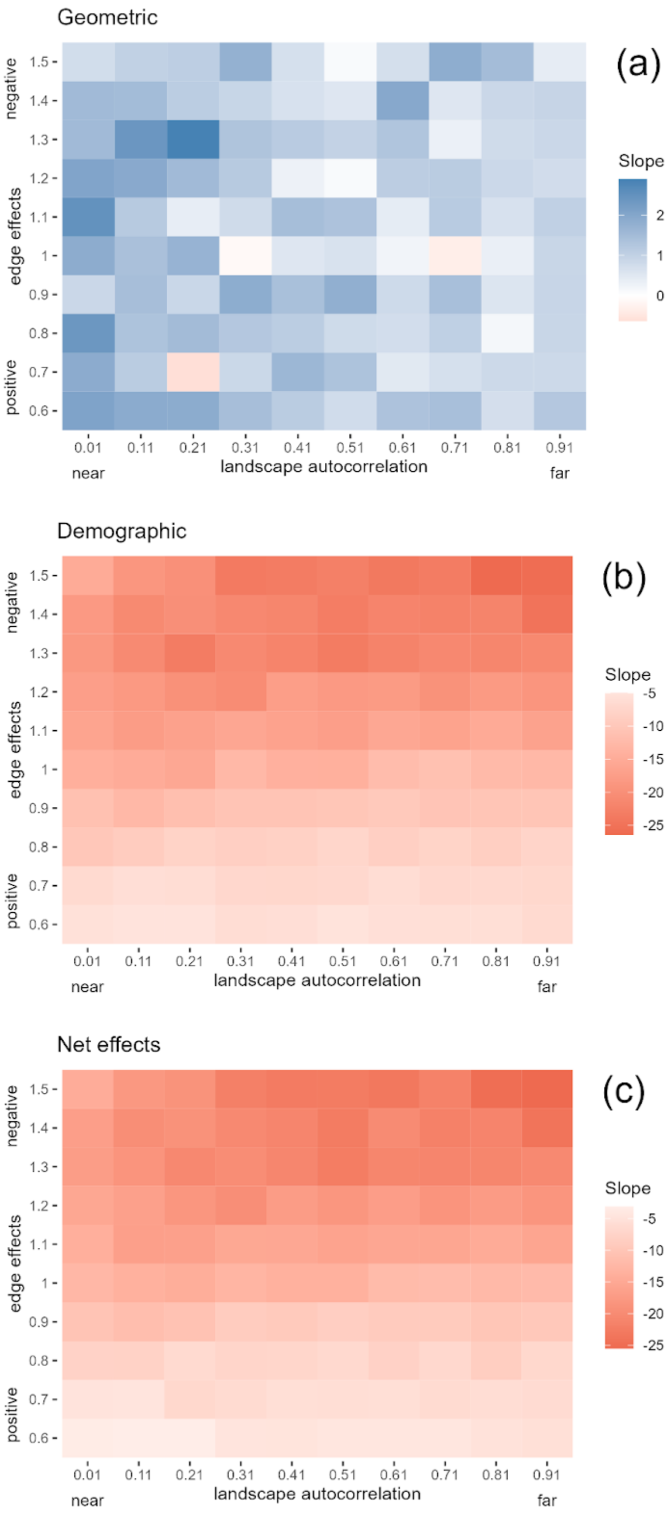


*Figure S3B: A full factorial design testing the interplay between fragmentation effects at the sample scale by varying the edge effects and landscape autocorrelation parameters. Plot colours represent the slope of a regression line for species richness with increasing levels of fragmentation (0.1-0.9) immediately after fragmentation for geometric effects (a) and at the end of the simulation for net effects (c). Demographic effects were calculated by deducting the geometric effects slope value from the net effect slope value (b). Slope results for each box were calculated from 9 simulations with varying fragmentation levels repeated 10 times. Values for species richness at the sample scale are based on mean values of 30 samples (single cells).*
